# Arbuscular mycorrhizal fungus *Rhizophagus irregularis* expresses an outwardly Shaker-like channel involved in rice potassium nutrition

**DOI:** 10.1101/2022.11.04.515200

**Authors:** Claire Corratgé-Faillie, Louise Matic, Layla Chmaiss, Houssein Zhour, Jean-Pierre Lolivier, Pierre-Alexandre Audebert, Xuan Thai Bui, Maguette Seck, Kawiporn Chinachanta, Cécile Fizames, Daniel Wipf, Hervé Sentenac, Anne-Aliénor Very, Pierre-Emmanuel Courty, Doan Trung Luu

## Abstract

Potassium (K^+^) plays crucial roles in many physiological, molecular and cellular processes in plants. Direct uptake of this nutrient by root cells has been extensively investigated, however, indirect uptake of K^+^ mediated by the interactions of the roots with fungi in the frame of a mutualistic symbiosis, also called mycorrhizal nutrient uptake pathway, is much less known. We identified an ion channel in the arbuscular mycorrhizal (AM) fungus *Rhizophagus irregularis*. This channel exhibits the canonical features of Shaker-like channel shared in other living kingdoms and is named RiSKC3. Transcriptionally expressed in hyphae and in arbuscules of colonized rice roots, RiSKC3 has been shown to be located in the plasma membrane. Voltage-clamp functional characterization in *Xenopus* oocytes revealed that RiSKC3 is endowed with outwardly-rectifying voltage-gated activity with a high selectivity for potassium over sodium ions. RiSKC3 may have a role in the AM K^+^ pathway for rice nutrition in normal and salt stress conditions. The current working model proposes that K^+^ ions taken up by peripheral hyphae of *R. irregularis* are secreted towards the host root into periarbuscular space by RiSKC3.

**Significance Statement:** Arbuscular mycorhizal fungus *Rhizophagus irregularis* expresses a Shaker-like channel, located in the plasma membrane, endowed with a strictly outwardly-rectifying voltage-gated activity with a high selectivity for potassium over sodium ions. The current working model proposes that K^+^ ions taken up by peripheral hyphae of *R. irregularis* are secreted towards the host root into periarbuscular space by this Shaker-like channel.

## Introduction

The low availability of nutrient ions in the soil is one of the main abiotic constraints limiting crop productivity. In conventional agriculture, this can be overcome by large inputs of fertilizers, but at high ecological (and economic) costs. Today, however, there is a strong demand for a new revolution in agriculture, aimed at reducing its environmental costs while coping with the world’s growing population (1, 2). Many fronts of research are opened likely to contribute to such a revolution, among which the analysis of the beneficial interactions between roots and mycorrhizal fungi, which improve plant mineral nutrition. The present study deals with arbuscular mycorrhization (AM) and potassium nutrition. AM symbiosis is thought to have played a crucial role in facilitating the colonization of land by early plants. It can be established by most plant species, including major crops. Potassium (K^+^) is a macro-nutrient playing very diverse and essential roles in plants. While it is required in large quantities, *e.g.* its concentration in the cytosol has to be close to 100 mM, its availability in most soils is relatively low in absence of fertilization, which limits plant growth and affects plant health. In the last report of the FAO on world fertilizer trends and its outlook, the demand for mineral fertilizer would has increased by a total of 9% between 2016 and 2022, and that for K^+^ by 14% (3).

Compared with the contribution of AM symbiosis to plant phosphate (P) and nitrogen (N) nutrition, the contribution to K^+^ nutrition has been found to be less evident (4) and even insignificant in some studies (5). However, substantial effects of AM symbiosis on plant K^+^ nutrition have also been observed (6–12). Interestingly, in soybean, large differences in growth response to AM symbiosis with different isolates of *Glomus mosseae* seemed to be more related to improved K^+^ rather than P nutrition of the host plant (13). Taken together, these reports suggest that the beneficial effect of AM symbiosis on plant nutrition may be more dependent on the fungal partner species/isolate and environmental conditions in the case of K^+^ nutrition than P or N nutrition.

The mycorrhizal pathway enabling nutrient ion translocation from the soil to the host plant roots via the fungus partner is also better characterized for P and N than for K^+^. This pathway comprises at least 3 essential membrane transport events, namely ion uptake by external hyphae exploring the soil and; after diffusion from external to internal hyphae, ion secretion by fungal cells present at the interface with the root, and finally uptake by root cells of the ions secreted by the fungus at this interface. No membrane transport system involved in fungal secretion to host root cells has yet been identified, whatever the nutrient. For P and N, molecular-level information is available on the other two essential transport events of the mycorrhizal pathway, i.e. uptake from the soil by external hyphae and, at the interface with the internal hyphae, re-absorption by root cells of nutrients secreted by the fungus, given that P, NO_3_^-^ and NH_4_^+^ transport systems involved in the corresponding uptake events are identified (14). For K^+^, no membrane transport system involved in any of the 3 essential transport events of the endomycorrhizal pathway was identified so far.

Here we investigate the beneficial effects on plant K^+^ nutrition of AM symbiosis between rice (cv Kitaake) and *Rhizophagus irregularis* (DAOM197198). A strongly significant effect of the symbiotic interaction on plant growth was observed but only under low K^+^ availability conditions, and especially in presence of high Na^+^ external concentrations, the latter stress being known to impede K^+^ uptake by roots. K^+^ (and rubidium (Rb^+^), used as a K^+^ tracer) assays revealed that the AM symbiosis strongly increased the plant capacity to take up K^+^ and the total K^+^ contents by ca. 100% in such conditions. A membrane transport system likely to be involved in K^+^ secretion towards the host plants, namely an outwardly rectifying channel from the Shaker K^+^ channel superfamily extensively studied in animals and plants, is characterized, revealing unique functional features, especially in terms of sensitivity to membrane voltage.

## Results

### Beneficial effects of inoculation with *R. irregularis* on rice growth and K^+^ nutrition

The effects of inoculation with *R. irregularis* on shoot growth and K^+^ nutrition were investigated in the short-live cycle rice cultivar Kitaake in controlled conditions. The plants were inoculated or not with the fungus and grown at two different K^+^ concentrations for 9 weeks in pots filled with sand to control the amount of nutrients provided. They were watered with Yoshida-modified-solution containing either 0.8 mM or 16-fold less K^+^ (0.05 mM), the corresponding solution being here qualified of “high” or “low” K^+^, respectively. The latter condition actually resulted in limited K^+^ availability (K^+^ shortage) since, in the absence of mycorrhization, the plants (NM plants, for non-mycorrhized) displayed significant reductions in shoot dry weight (about 27% decrease), K^+^ content (∼73% decrease) and total K^+^ amount (∼80% decrease) when compared with NM plants supplied with normal K^+^ solution (Fig. 1). Interestingly, in such conditions of low K^+^ availability, the interaction with *R. irregularis* was found to significantly improve plant K^+^ nutrition and growth: AM plants exhibited an increase in shoot dry weight, K^+^ content and total K^+^ amount, when compared with the NM plants, by ∼21%, ∼91%, and ∼137%, respectively (Fig. 1). On the other hand, when the plants were grown under the high K^+^ availability regime, inoculation with *R. irregularis* was without any significant effect on plant growth or K^+^ nutrition (Fig. 1).

**Figure 1.**
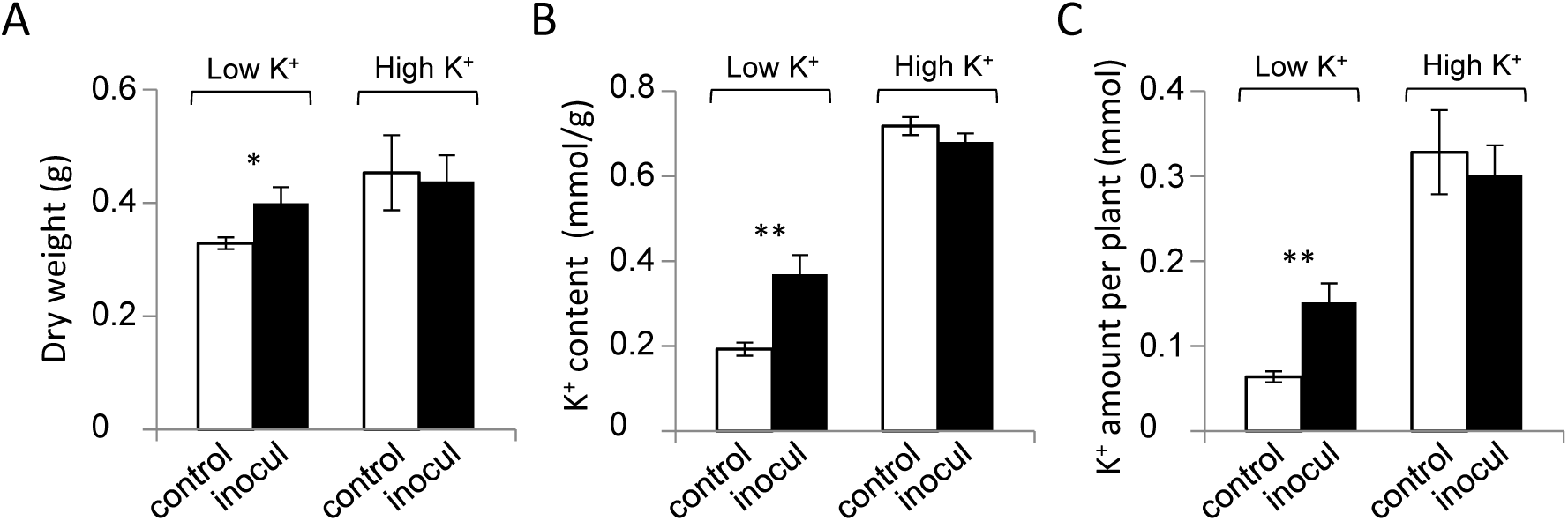
Effects of *R. irregularis* inoculation on *O. sativa* growth and K^+^ nutrition. Rice seedlings germinated for one week on water were transplanted in sterilized quartz sand in pots inoculated (inocul) or not inoculated (control) with *R. irregularis*. They were grown in green house for 9 weeks, being watered with a nutrient solution containing either 0.05 mM K^+^ or 0.8 mM K^+^ (low K^+^ and high K^+^ treatment, respectively). They were then harvested for measurement of shoot dry weight (**A**), K^+^ content (mmol/g DW, **B**) and total K^+^ amount (mmol per plant, **C**). Mean values and SEMs for 10 plants from two independent experiments. Significant differences (Student’s *t*-test) between the plants grown in the control or inoculated conditions are indicated: ** for *P*<0.001, * for *P*<0.05.

Colonization of the roots by the fungus at the end of the 9-week growth period was monitored in the two types of plants, grown under low or high K^+^ availability. Trypan blue staining of root samples revealed no significant difference between the high– and low-K^+^ conditions*, i.e.* with a mycorrhization frequency of ∼80%, an intensity of the mycorrhizal colonization of ∼25% and an arbuscule abundance of ∼10% whatever the availability of K^+^, low or high, during plant growth (Supplemental Fig.1). Thus the absence of beneficial effects of the fungus in plants grown at high K^+^ availability did not result from the absence of mycorrhization in these conditions but may be hypothesized to result from the fact that, in such conditions of K^+^ availability, the plant is no longer dependent on its fungal partner for its K^+^ nutrition. It is however worth to note that the corresponding plants watered with the high K^+^ solution were engaged in the symbiotic interaction with the fungus although they apparently did not get any benefit from the symbiosis in such conditions of high K^+^ availability.

### Beneficial effects of *R. irregularis* inoculation on rice tolerance to salinity

Since plant tolerance to soil salinity is crucially dependent on K^+^ homeostasis in shoots, we investigated the effects of *R. irregularis* inoculation on Kitaake rice tolerance to saline conditions. A salt stress was applied for a week to NM and AM plants grown for 8 weeks in sand. The salt stress solution contained 0.8 or 0.05 mM K^+^ (high or low K^+^ condition, respectively) and the commonly used K^+^ tracer Rb^+^ (12, 15, 16) at the same concentration as K^+^. As in the previous experiment (Fig. 1), differences between AM and NM plants were observed only in the low K^+^ availability regime. At the end of the one-week salt stress treatment, the AM plants grown under the low K^+^ regime again showed higher shoot dry weight, K^+^ content and K^+^ amount than the corresponding NM plants, by ∼32%, ∼110% and ∼179% respectively (Fig. 2A-C). The shoot contents in Rb^+^ were also found to be very significantly higher, by ca. 2 times, in the AM than NM plants (Fig. 2D). This indicates that the differences in K^+^ contents and amounts observed at the end of the salt stress period (Fig. 2B-C) cannot be merely ascribed to differences established during the previous 8 weeks of plant growth in the absence of salt treatment but that they are a consequence, at least in part, of much stronger capacity to take up K^+^ in presence of high Na^+^ concentrations, during the salt treatment itself, in AM than NM plants. Altogether, these results provide evidence that endomycorrhizal symbiosis with *R. irregularis* favours K^+^ uptake and accumulation in presence of stressing Na^+^ concentrations under low K^+^ availability conditions.

**Figure 2.**
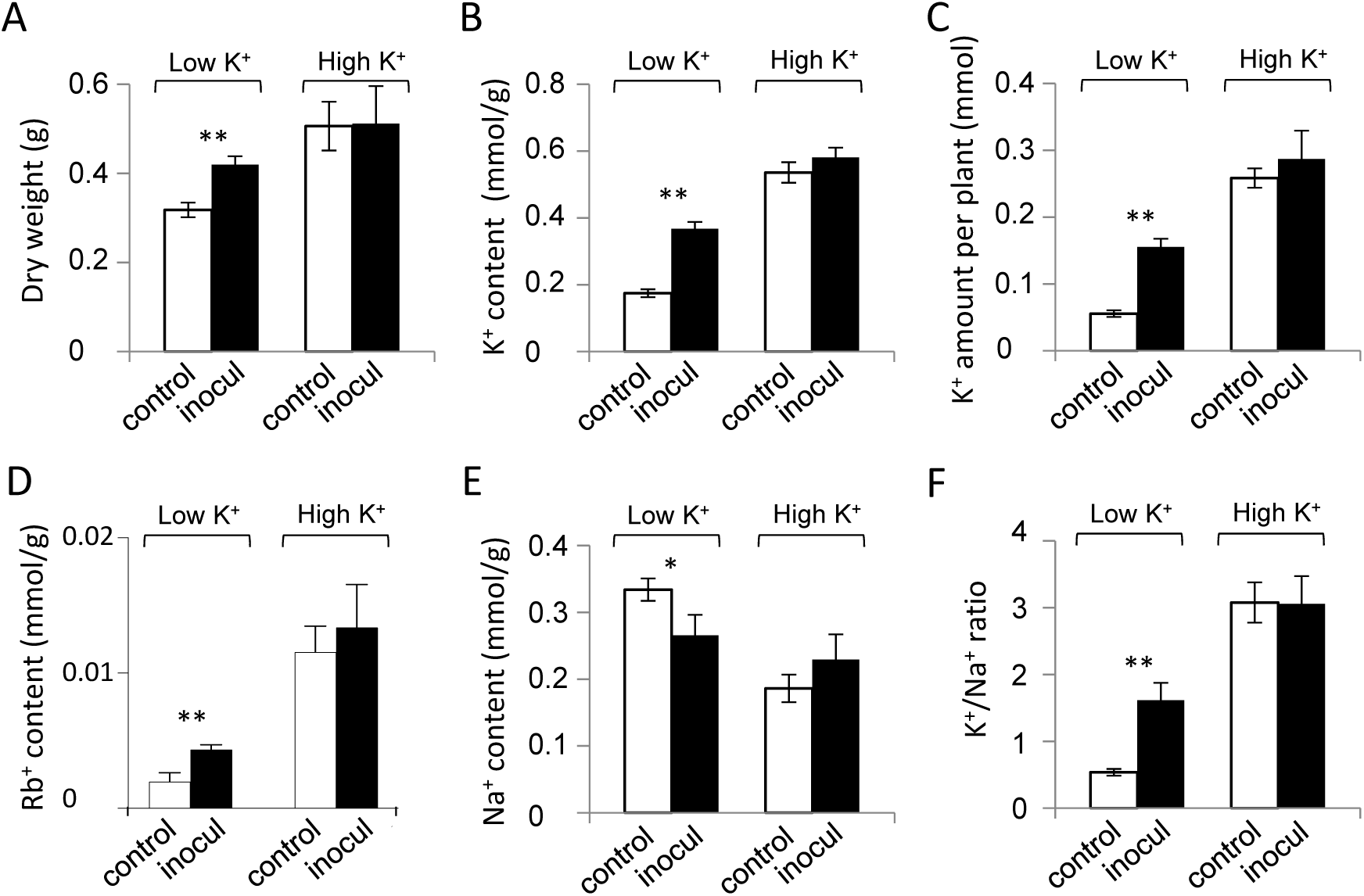
*R. irregularis* inoculation improves rice tolerance to saline conditions. Rice seedlings germinated for one week on water were thereafter grown in sand inoculated or not inoculated with *R. irregularis* (inocul and control treatment, respectively). They were watered with a nutrient solution containing either 0.05 mM K^+^ or 0.8 mM K^+^ (low K^+^ and high K^+^ treatment, respectively) for 8 weeks. They were then subjected to salt stress conditions by adding 100 mM NaCl to the watering solutions. Rb^+^ (used as a classical K^+^ tracer) was also added (as RbCl) into both nutrient solutions at the same concentration (0.05 or 0.8 mM) as K^+^. The plants were then grown for one further week, being watered with the same solutions but free from added NaCl and RbCl, before being harvested for measurement of shoot dry weight (g, **A**), K^+^ content (mmol/g DW, **B**), K^+^ amount per plant (mmol, **C**), Rb^+^ content (mmol/g DW, **D**), Na^+^ content (mmol/g DW, **E**) and K^+^/Na^+^ content ratio (**F**). Mean values and SEMs from two independent experiments, with a total number of plants of ten. Significant differences (Student’s *t*-test) between the plants grown in the control or inoculated conditions are indicated: ** for *P*<0.001, * for *P*<0.05.

Plant capacity to maintain low Na^+^ contents in shoots, and consequently high K^+^/Na^+^ shoot content ratios, is a major determinant of tolerance to salinity in non-halophyte species like rice. The saline treatment resulted in large accumulation of Na^+^ in shoots of all types of plants. In plants grown under the high K^+^ availability regime, the endomycorrhizal interaction with *R. irregulari*s was without any significant effect on Na^+^ shoot contents and K^+^/Na^+^ shoot content ratio (Fig. 2E-F). In contrast, in plants grown under the low K^+^ availability condition, the interaction with *R. irregularis* led to a significant reduction in shoot Na^+^ contents, by about 20% (Fig. 2D). This and the increase in K^+^ contents due to mycorrhization (Fig. 2C) resulted in a strong increase in the K^+^/Na^+^ shoot content ratio, by about 300% when compared with the corresponding NM plants (Fig. 2E). Combined with the positive effect on plant growth (Fig. 2A), these results reveal that plant tolerance to salt stress conditions in presence of reduced K^+^ concentration can be significantly increased in rice by endomycorrhizal symbiosis with *R. irregularis*.

Characterization of one of the 4 K^+^ transport systems identified in *R. irregularis* (17, 18), RiSKC3, was conducted to shed light on the endomycorrhizal pathway of K^+^ transport to the host plant.

### RiSKC3 exhibits canonical features of K^+^ channels from the Shaker superfamily

Sequence analysis indicates that RiSKC3 shares strong homologies with K^+^ channels belonging to the Shaker superfamily. Based on the present knowledge, Shaker K^+^ channels are present in all animals and plants (19–22). Searches in genomic data bases indicate that they are also present in different fungi, but not in all. For instance, Shaker K^+^ channels can be identified in the model ectomycorrhizal species *Hebeloma cylindrosporum* and *Laccaria bicolor*, but not in the model yeast *Saccharomyces cerevisiae* nor in almost all of the endophytic fungi whose genome sequences are present in databases ((23), and our own searches in JGI (24)).

Shaker channels are tetrameric proteins build of 4 polypeptides encoded by Shaker genes. A Shaker polypeptide typically displays a transmembrane hydrophobic core comprised of 6 transmembrane domains, named S1 to S6, and a pore-forming domain (P) present between S5 and S6 and playing a determinant role in the building and functioning of the channel selectivity filter (21, 25). A schematic comparison of the predicted secondary structure of RiSKC3, where such a hydrophobic core can be identified, with that of three outwardly-rectifying K^+^ channels (see below: RISK3 functional characterization) from the same “Shaker-like” superfamily, the human HsKcnh1 K^+^ channel (26)), the *Drosophila melanogaster* Shaker DmShB (27) and the *Arabidopsis thaliana* AtSKOR K^+^ channel (28) is shown in Figure 3A, drawn to scale using the first amino acid of the predicted S1 domain to start the alignment. In Shaker channels, the S4 transmembrane segment comprises positively charged residues (presence of arginine, lysine and histidine amino acids), which enables this segment to act as a voltage sensor: S4 movements within the membrane in response to changes in electrical polarization of the membrane result in channel activation or inactivation (29). The S4 segment of RiSKC3 harbors 4 arginine (R) and 1 lysine (K) residues (R265, R271, R274, K280, R290), the 4 R being present at strictly conserved positions when compared with DmShB and AtSKOR (Fig. 3B). Finally, the pore domain identified in RiSKC3 harbors a GYGD motif (Fig. 3B), which is a hallmark determinant of the selectivity for K^+^ in ion channels (29, 30).

**Figure 3.**
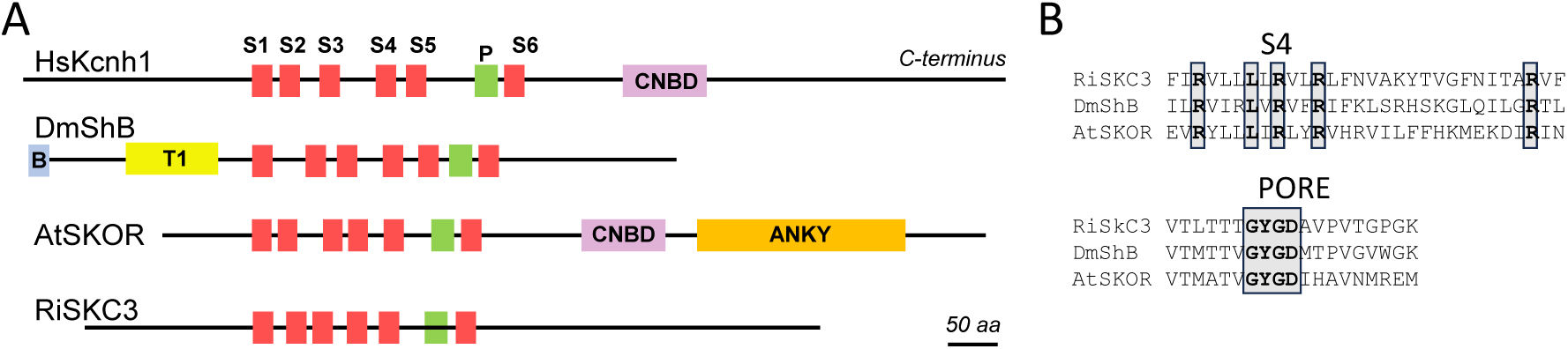
Analysis of RiSKC3 secondary structure. (**A**) RiSKC3 sequence was compared to the following sequences: HsKcnh1 (an inward Shaker-like K^+^ channel from *Homo sapiens*, taxid:9606), DmShB (an outward Shaker from *Drosophila melanogaster,* taxid:7227) and AtSKOR (an outward Shaker from *Arabidopsis thaliana* taxid:3702). The schematic secondary structure of these 4 channel polypeptides is shown drawn to scale: the length of the black lines corresponds to the length of the channel polypeptides, and the length and position of the different domains are displayed according to this scale. The 4 polypeptides are aligned with respect of the first amino acid of the predicted S1 domain. All Shaker channels typically display a hydrophobic core consisting of 6 transmembrane segments (named S1 to S6, in red) and a pore domain P (in green) between S5 and S6. The N– and C-terminal regions upstream and downstream the transmembrane hydrophobic core are on the cytosolic face of the membrane. Different domains have been identified in these regions, among which a Ball-and-Chain domain (B, in blue) and a Tetramerization domain (T1, yellow) in DmShB, a Cyclic Nucleotide-Binding Domain (CNBD, pink) in HsKcnh1 and AtSKOR (and in every Shaker channel from plants, where this domain has been shown to be involved in channel tetramerization), and an Ankyrin domain (ANKY, orange) in AtSKOR and many other Shaker channels from plants. (**B**) S4 segment and Pore domain (upper and lower panel, respectively) of RiSKC3 aligned with the corresponding sequences in DmShB and AtSKOR. The protein (GenBank) accession numbers are provided in Supplemental Table S1.

It is however interesting to note that the RiSKC3 regions present upstream of S1 and downstream of S6, both of which have been shown to be present on the cytosolic face of the membrane in Shaker channels (29), do not display any of the functional domain presently identified in K^+^ channels from the Shaker superfamily, such as the so-called ball-and-chain domain (B in Fig. 3A) and the Tetramerization domain (31, 32) (T1 in Fig.3A) identified in DmShB, the cyclic-nucleotide-binding (homology) domain (CNBD in Fig. 3A) identified in HsKcnh1 (26) and in AtSKOR (28) or the ankyrin domain (ANKY in Fig. 3A) identified in AtSKOR and other plant Shaker-like K^+^ channels (33, 34). CNB(H)D domains have been shown to contribute to channel tetramerization and regulation of channel activity (35, 36) and ANKY domains constitutes a site of interaction with regulatory proteins (e.g., kinases or phosphatases) controlling channel activity (37, 38). RiSKC3 should thus contain functional domains, e.g. involved in channel tetramerization and regulation, which have not been identified by the present *in silico* searches and might be specific of Shaker channels in (endomycorrhizal) fungi.

The phylogenetic relationships of RiSKC3 with the DmShB, AtSKOR, HsKcnh1 channels evoked above, with the Shakers HcSKC and LbSKC from the ectomycorrhizal fungi *Hebeloma cylindrosporum* and *Laccaria bicolor*, and with 3 other members from the Shaker super-family, AtAKT1 from *Arabidopsis thaliana*, PpAKT1 from *Physcomitrella patens* and the human Shaker HsKv4.2 have been analyzed (Supplemental Fig. 2). RiSKC3 appears to be closer from the fly DmShB and human HsKv4.2 Shakers and from the two fungal Shaker proteins than from the selected plant K^+^ channels and the human HsKcnh1.

### *RiSKC3* transcripts are localized in hyphae and arbuscules

As a first step, the localization of *RiSKC3* transcripts was studied in mycorrhizal rice roots (Figure 4). On the plant side, RT-PCR allowed amplification of the rice housekeeping gene *ubiquitin-like protein SMT3* in both NM and AM conditions (Fig. 4A). As a positive control, the phosphate transporter transcript *OsPT11* (AF536960), which is known to be specifically expressed in AM condition (39), was consistently detected only in AM condition (Fig. 4A). On the fungal side, the housekeeping gene *cyclophilin-like domain-containing protein RiCYP1* was detected only in AM condition. Regarding *RiSKC3*, RT-PCR detected transcripts from this gene only in AM condition (Fig. 4A). Relative quantification by RT-qPCR revealed that *RiSKC3* transcripts accumulated in roughly identical proportions under the conditions of low or high external K^+^ availability. (Fig. 4B).

**Figure 4.**
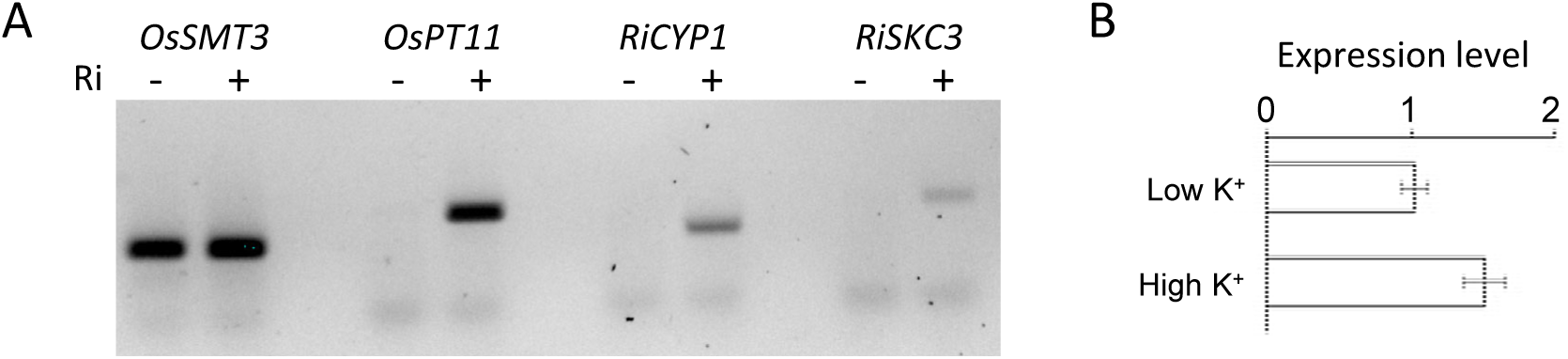
Expression of *RiSKC3* in AM rice roots. Rice plants were grown for 9 weeks in sand inoculated (+) or non-inoculated (−) with *R. irregularis* as described in legend to Figure 1. Roots were then collected and total RNA extracted. (**A**) RT-PCR analysis of *RISKC3* expression in roots from plants watered with low K^+^ solution. RT-PCR was performed to amplify ∼100 bp fragments from the coding sequence of either *OsSMT3*, *OsPT11*, *RiCYP1* or *RiSKC3* genes. (**B**) RT-qPCR comparison of *RiSKC3* expression in plants watered with the nutrient solution containing either 0.05 mM or 0.8 mM K^+^. Means and SEMs from two independent experiments.

The cellular localization of *RISKC3* transcripts was then analysed by means of *in situ* RT-PCR experiments to take advantage of the high sensitivity of this technique in detecting low-abundance transcripts (Fig. 5). It is worth to note that endogenous alkaline phosphatase activity was detected in fixed tissues (Supplemental Fig.3). However, when the tissues underwent the PCR cycles at maximum 95°C in the thermocycler, this endogenous activity vanished, then allowing detection of transcripts by means of exogenous alkaline phosphatase activity (Fig. 5A-B). Using this procedure, transcripts of the rice gene *OsPT11* were detected in root cells colonized by an arbuscule but not in hyphae (Fig. 5C-D), in agreement with previous analyses (39). Regarding *RiSKC3* expression, transcripts from this gene were detected in arbuscules and hyphae (Fig. 5E-F).

**Figure 5.**
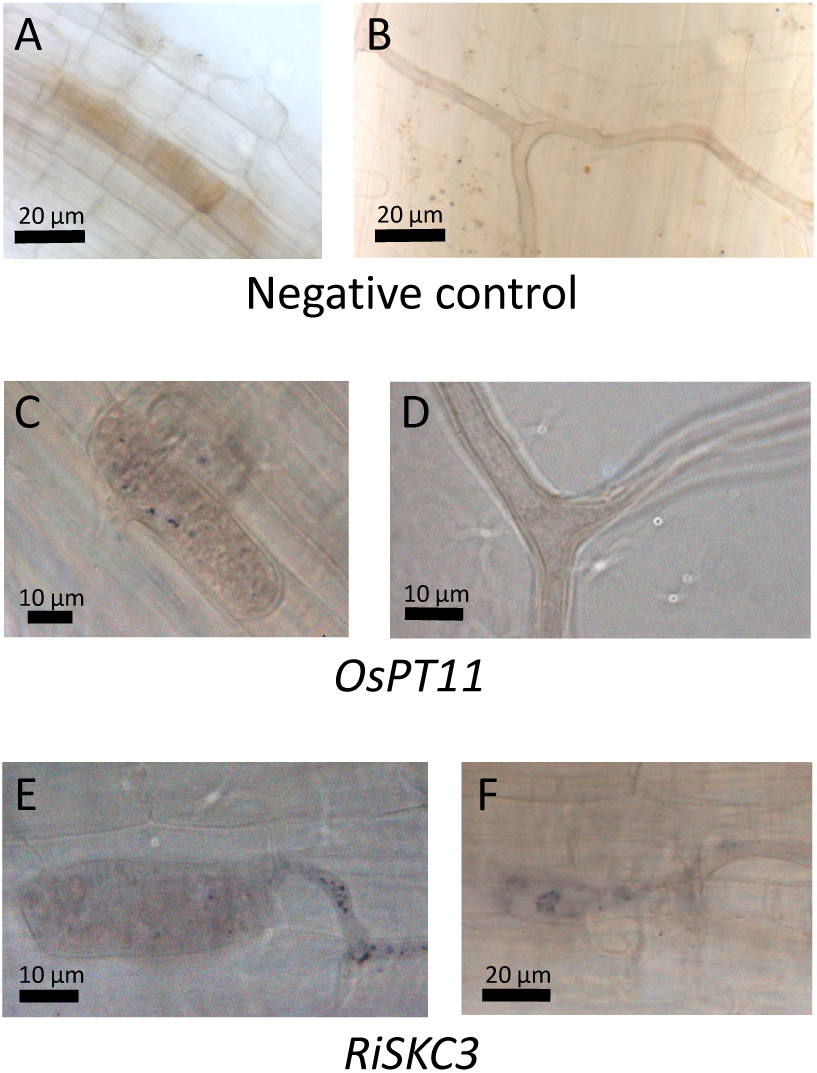
Expression pattern of *RiSKC3* in AM rice roots. Rice plants inoculated with *R. irregularis* were grown for 9 weeks in sand watered with low K^+^ solution before root samples were collected for whole-mount *in situ* RT-PCR experiments. As negative control, fixed tissues were subjected to PCR cycles and incubated with reagent for labeling and with NBT/BCIP. No phosphatase-alkaline activity was detected in cells colonized by arbuscules (**A**) or in extraradical hyphae (**B**). Using primers specific to *OsPT11*, *in situ* RT-PCR was performed and signal was detected in root cells colonized by arbuscules (**C**), but not in extraradical hyphae (**D**). Using primers specific to *RiSKC3 l*ed to detection of signal in root cells colonized by arbuscules (**E**) and in intraradical hyphae (**F**).

### RiSKC3 is a plasma membrane-localized protein

The subcellular localization of RISKC3 was investigated by means of heterologous expression in *S. cerevisiae.* The coding sequence of RISKC3 was fused in frame with the eGFP reporter gene downstream of the *S. cerevisiae phosphoglycerate kinase* (*PGK*) promoter in the pFL61 expression vector, and the construct was expressed in yeast. Samples were stained using FM4-64, a plasma membrane-specific dye. Confocal microscopy showed a colocalization of the GFP and FM4-64 fluorescence in the cells expressing the RISKC3-eGFP construct (Fig. 6). This result, together with the data obtained by heterologous expression of RISKC3 in *Xenopus* oocytes, revealing exogenous K^+^ channel activity at the cell membrane (see below), provide evidence that *RiSKC3* encodes a plasma membrane-localized K^+^ channel.

**Figure 6.**
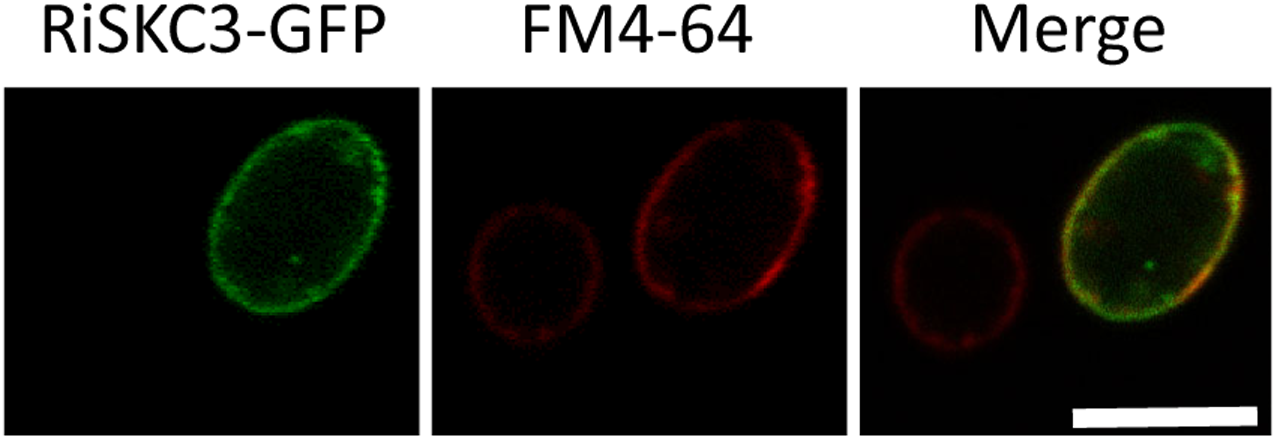
Subcellular localization of RiSKC3. Yeast cells were transformed with a *RiSKC3-GFP* reporter construct. Confocal microscopy analyses were carried out after staining with membrane-dye FM4-64. Representative image of a yeast cell under GFP channel (left panel) or FM4-64 channel (middle panel), and merged image (right panel).

### Functional characterization in *Xenopus* oocytes

The functional properties of RiSKC3 were investigated by carrying out electrophysiological analyses after heterologous expression in Xenopus oocytes (Fig. 7). Using the two electrode voltage-clamp technique, we observed that RiSCK3 expression resulted in an outwardly-rectifying conductance in oocytes injected with *RiSKC3* cRNA. Typical current recordings obtained in control oocytes injected with water and in RiSKC3-expressing oocyte upon depolarization of the membrane in external medium containing 10 mM K^+^ are shown in Figure 7A. The macroscopic current displayed a slow activation upon membrane depolarization, no inactivation and a slow deactivation upon membrane repolarization. Increasing the external K^+^ concentration from 10 to 100 mM resulted in a decrease in K^+^ outward current as shown on I–V (current-voltage) curves (Fig. 7B), which was consistent with the hypothesis that RiSKC3 is permeable to K^+^. Tail-current analysis was then performed (insets in Figure 7C) to determine the shift in current reversal potential (Erev) upon the replacement of 100 mM K^+^ in external medium by 10 mM K^+^. This 10-fold decrease in external K^+^ concentration resulted in a –51 mV shift of Erev (Fig. 7C). The magnitude of this shift, which is close to the predicted theoretical shift of Erev in a channel that would be exclusively permeable to K^+^, –58 mV, thus indicated that RiSKC3 displays a relatively high selectivity for K^+^.

**Figure 7.**
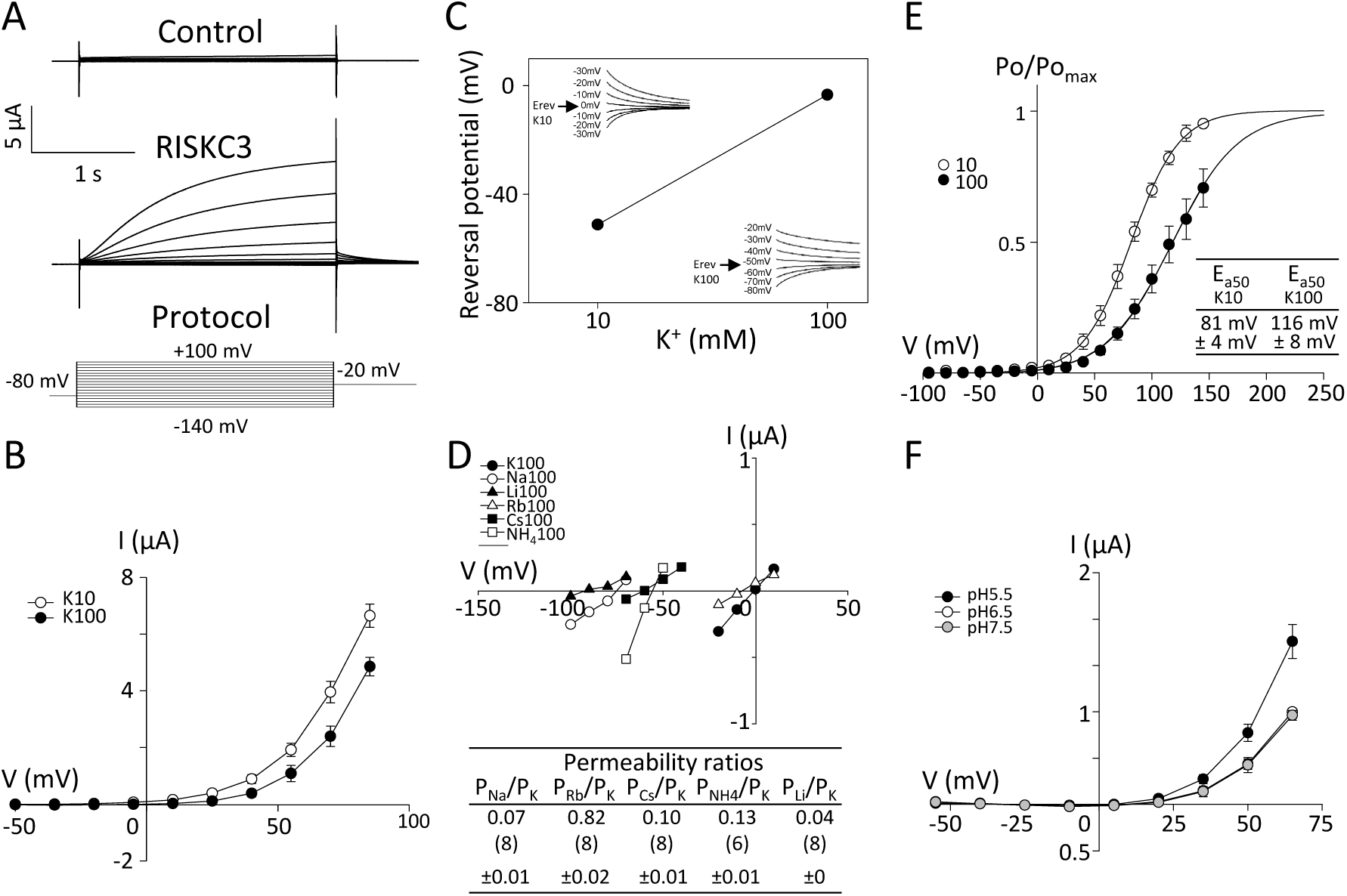
Functional characterization of RiSKC3 in *Xenopus* oocytes. (**A**) Typical current traces recorded in 10 mM KCl in control oocytes (injected with water) or in oocytes injected with *RiSKC3* cRNA using a voltage clamp protocol (bottom panel) with voltage clamp pulses ranging from −140 to + 100 mV, in increments of 15 mV from a holding potential of –80 mV. (**B**) RiSKC3 current–voltage relationships in 10 mM (white circles) or 100 mM KCl (black circles). Mean value ± SE, from n=4 oocytes. (**C**) Variation of RiSKC3 reversal potential (Erev) in 10 mM and 100 mM KCl. Erev was determined using a tail-current protocol: after activation of RiSKC3 at +55 mV, voltage pulses were performed at voltages flanking Erev. Examples of Erev measurement are shown for 10 mM KCl (upper left inset) and for 100 mM KCl (lower right inset). (**D**) Permeability of RiSKC3 to different monovalent cations. Representative current–voltage relationships of RISKC3 deactivation currents, which were recorded using a tail-current protocol (as described above) in 100 mM of either K^+^, Na^+^, Li^+^, Rb^+^, Cs^+^ or NH4^+^. Permeability ratios of the different cations with respect to that of K^+^ were calculated from the variations of Erev using the Goldman–Hodgkin–Katz equation. Ratios are given as mean ± SE (number of independently obtained values provided between parenthesis). (**E**) Effect of the membrane voltage and the external K^+^ concentration on RiSKC3 open probability. The relative open probability (Po/Pomax) was obtained from the analysis of deactivation currents upon return to the holding potential (Lebaudy *et al*., 2010; mean ± SE, n=5). The solid lines are Boltzmann fits to the mean Po/Pomax values. The mean values (± SE, n=5) of the half-activation potential (Ea50) of RiSKC3 obtained from these fits in 10 mM or 100 mM of KCl are provided in the inset. (**F**) Activation of RiSCK3 currents by external acidification. The external solution contained 100 mM KCl at pH 5.5, 6.5 or 7.5. Means ± SE, n = 4. The bath solution contained 1 mM CaCl2, 1.5 mM MgCl2, 10 mM HEPES, pH 7.5 (or 10 mM MES for pH 5.5 and 5 mM MES 5 mM HEPES for pH 6.5) as standard background medium, and 10 mM KCl or 100 mM KCl, or 100 mM NaCl, LiCl, RbCl, CsCl or NH4Cl.

The ionic selectivity of RiSKC3 was then further investigated by replacing K^+^ in the external medium by several alkali cations (Na^+^, Li^+^, Rb^+^, Cs^+^ or NH_4_^+^) at the same 100 mM concentration. By determining the current reversal potential (Erev) from classical tail-current recording analyses, relative permeability ratios (to K^+^) were calculated using the Goldman-Hodgkin-Katz equation, which revealed the following permeability order: K^+^>Rb^+^>>NH_4_^+^>Cs^+^>Na^+^>Li^+^ (Fig. 7D bottom panel). The relative permeability of RiSCK3 to Rb^+^ is high (0.82), thus indicating that this cation can be considered as a good tracer of K^+^ with respect to RiSKC3 activity. Conversely, the low value of RiSKC3’s relative permeability to Na^+^, combined with the fact that the concentration of K^+^ in the fungal cytosol is probably high, in the 100 mM range, as in all living cells, and that the cytosolic concentration of Na^+^ is probably lower, strongly suggests that the flow of cations secreted into the external environment through RiSKC3 consists essentially of K^+^ ions.

The current-voltage curves shown in Figure 7B revealed that RiSKC3 activation occurred at very depolarized membrane potential, *e.g.* close to 0 mV in presence of 10 mM K^+^ in the external medium. For comparison, outwardly-rectifying Shaker channels characterized in plants display activation potentials close to –50 mV in similar external conditions (10 mM K^+,^ pH ∼7.5). The voltage sensitivity of RiSKC3 activation was further investigated by recording deactivation currents to determine the channel relative open probability (Po/Pomax) in the presence of 10 or 100 mM external K^+^. The experimental curves describing the voltage-dependency of the channel relative open probability were fitted with the classical two-state Boltzmann law as displayed in Figure 7E. The results indicate that the channel half-activation potential (Ea50, the membrane potential at which the channel relative open-probability is 0.5) is close to +80 mV in 10 mM external K^+^, For comparison, the corresponding value for the plant outward Shaker SKOR is close to 0 mV (40). The data displayed by Figure 7E also indicate that the RiSKC3 half-activation potential is dependent on the external concentration of K^+^, being shifted by ca. +35 mV upon a 10-fold increase of this concentration, from 10 to 100 mM (Fig. 7E). A shift in half-activation of similar magnitude upon the same increase in external K^+^ concentration (from 10 mM to 100 mM) has been reported in the Arabidopsis AtSKOR channel. The gating of RiSKC3 and that of AtSKOR are thus sensitive both to membrane voltage and to the extracellular concentration of K^+^. Such a double regulation has been proposed in AtSKOR to ensure that the channels open only when the driving force for net K^+^ flux is directed outwards (40, 41). Endowed with such a double regulation, and the fact that the former regulation by voltage opens the channel only at very depolarized membrane potentials, RiSKC3 can be hypothesized to be absolutely strictly dedicated to K^+^ secretion from the cell.

Finally, a series of experiments were carried out to determine the pH sensitivity of RiSKC3. Comparison of the I–V curves obtained at pH 5.5, 6.5 and 7.5 in the presence of 100 mM K^+^ in the external medium revealed that RiSKC3 is activated by the acidification of the external medium, being increased by 170% at pH 5.5 (Fig. 7F).

## Discussion

### The rice-*R. irregularis* couple provides an appropriate model for studying the mechanisms by which AM symbiosis improves potassium nutrition

Root system interactions with mycorrhizal or endophytic fungi benefit the plant mineral nutrition. Essentially, the plant gives carbon compounds to its fungal partners, who in turn transfer nutrient ions taken up from the soil to the plant (14, 42, 43). The interactions with endophytic fungal species have been discovered relatively recently and are still poorly characterized (23). Mycorrhizal symbioses are better known. Endomycorrhizal symbiosis is considered the oldest root symbiosis in evolution. Its appearance accompanied the emergence of the first terrestrial plants from aquatic environments. The ability to engage in this symbiotic association is present in most plants, including crops.

The positive effects of AM symbiosis on plant mineral nutrition are well documented. With respect to the 3 main macro-nutrients, evidence is available that AM symbiosis can be the main source of P for the plant, and can also make a significant contribution to N nutrition. The effects on K^+^ nutrition are less well documented and sometimes appear contradictory. It has been proposed that the effects of the symbiotic interaction on plant K^+^ nutrition depend largely on the species, or even the cultivar or ecotype, of the partner plant and fungus (7, 9, 10, 13). Our results indicate that rice (Kitaake cultivar) shows a very significant improvement in K^+^ nutrition and growth when mycorrhized with *R. irregularis* (DAOM197198). This symbiotic interaction increases the amount of K^+^ present in the plant by 137% (Fig. 1C), and promotes plant growth by 21% (Fig. 1A). It has been widely demonstrated that maintaining a high K^+^/Na^+^ ratio in plant tissues is an essential process in the plant ability to tolerate saline conditions. The rice-*R. irregularis* symbiont couple is therefore a very interesting model for analyzing the mechanisms involved in improvement by the fungus of the plant mineral nutrition, and in particular of K^+^ nutrition under saline conditions, as rice is a cereal of prime importance worldwide, and production of this plant is facing increasingly serious salinity problems due to climate change, especially salinization of the water in delta areas where it is grown.

### A role for fungal outward Shaker channels in the pathway of K^+^ translocation towards the host plant in endomycorrhizal symbiosis

The set of K^+^ transport systems appears to be more restricted in *R. irregularis* than in the model ectomycorrhizal fungi *Hebeloma cylindrosporum*. and *Laccaria bicolor.* Based on homology searches with animal, plant and bacterial K^+^ channels or transporters, the model ectomycorrhizal fungus *H. cylindrosporum* possesses 7 K^+^ transport systems, belonging to 4 different families: 1 K^+^ channel from the Shaker family, 1 K^+^ transporter from the HAK/KT/KUP family, 2 K^+^ transporters from the TrK family (the corresponding family being named HKT in plants), and 3 K^+^ channels from the TOK family. Based on similar sequence homology analyses, *R. irregularis* possesses 4 K^+^ transport systems in total, limited to 2 different families: the Shaker family, which comprises 3 members named RISKC1 to RISKC3, and the HAK/KUP/KT family with one member, named RiHAK (17, 18). The fungal pathway of mycorrhizal K^+^ transport is thus likely to be less complex to decipher in the endo-than in the ectomycorrhizal fungus.

The information on the functional properties of K^+^ transport systems of mycorrhizal fungi is still scant. No information at all is available in AM fungi. In ectomycorrhizal fungi, the available information is basically restricted to members from the TRK and TOK families of the fungus model *H. cylindrosporum* (44, 45). Based on analyses in heterologous systems (two-electrode voltage clamp in *Xenopus* oocytes and/or complementation of yeast mutant strains defective in K^+^ transport), HcTRK1 has been shown to mediate inwardly rectifying K^+^ (and Na^+^) currents, and HcTOK1 and HcTOK2.1 to mediate outwardly rectifying K^+^ currents. The present hypotheses, supported by gene expression localization analyses, is that HcTRK1 mediates K^+^ uptake from the soil by external hyphae, and HcTOK2.2 plays a role in K^+^ secretion towards the plant by internal hyphae (23, 44, 45). It has been reported that attempts to characterize the unique member of the *H. cylindrosporum* Shaker channel by heterologous expression in Xenopus oocytes or in yeast have failed (46), and no information has been published on the unique member of the HAK/KT/KUP family from this ectomycorrhizal fungus model. Thus, to our knowledge, the *R. irregularis* K^+^ transport systems, which belongs to the Shaker or the HAK/KT/KUP families, have no counterparts characterized at the functional level in ectomycorrhizal fungi, either in *H. cylindrosporum* or in other fungal species.

*In silico* analyses have concluded that most species of non-mycorrhizal endophytic fungi do not possess K^+^ channels of the Shaker family (23), which may indicate that the presence of such channels in the fungal partner is important for the establishment of beneficial mycorrhizal-style interactions between roots and fungi. Here, the functional properties of a Shaker K^+^ channel from *R. irregularis* are analysed using the two-electrode voltage clamp procedure following heterologous expression in Xenopus oocytes. This channel, named RiSKC3 in genome data bases (17), displays the classical 6-transmembrane hydrophobic core of the channels from this family (Fig. 3A), including the fourth transmembrane segment characterized by the presence of positively charged residues (Fig. 3B), which can be assumed to confer the role of voltage-sensor to this segment, and the so-called pore-forming region, between the 5^th^ and the 6^th^ transmembrane segments, harboring the hallmark motif GYGD of K^+^ selective channels. It is worth to note that the regions upstream and downstream of the transmembrane hydrophobic core in RiSKC3 do not display any of the typical domains identified in animal or plant Shaker channels and shown to play a role in regulation of channel properties, such as the ball-and-chain domain and the tetramerization domain found in *Drosophila* Shaker channels (31), or the CNBD found in some animal K^+^ channels like the human HsKcnh1 (26, 36) and in all plant channels from the Shaker-like family (21, 34), or the ankyrin domain present in many plant Shaker channels (20) (Fig. 3). Thus, RiSKC3 can be considered as displaying an original Shaker structure. For instance, in the absence in RiSKC3 of the so-called tetramerization domain and CNBD, the regions that are responsible for the tetramerization of the 4 *RiSKC3*-encoded polypeptide chains building the functional RiSKC3 protein would deserve to be identified.

Expressed in oocytes, RiSKC3 behaves as a K^+^-selective voltage-gated outwardly rectifying K^+^ channel (Fig. 7A-D). The very strong outward rectification indicates that the function of this channel is exclusively to mediate K^+^ efflux, with no significant contribution to K^+^ uptake even when the transmembrane K^+^ electrochemical potential is steeply inwardly directed*, e.g.* when the electrical potential difference is of –100 mV inside and the external concentration of K^+^ is 100 mM (Fig. 7E). It is thus the first membrane transport system identified as strictly involved in nutrient ion secretion by endomycorrhizal species. Expressed at the interface with the host plant (Fig. 5), in internal hyphae and arbuscules, RiSKC3 can thus be part of the mycorrhizal K^+^ pathway ensuring K^+^ secretion towards plant K^+^ uptake systems. Acidification of the plant-fungus interface, resulting from the activity of proton pump ATPases would stimulate K^+^ secretion towards the host plant due to the sensitivity of RiSKC3 to pH (Fig. 7F). The very positive values of its activation potentials (Ea50 found to be close to +80 mV in 10 mM K^+^ and +120 mV in 100 mM K^+^) make this channel truly remarkable compared with other outwardly-rectifying Shaker channels characterized in plants (e.g. AtSKOR Ea50 close to 0 mV in 10 mM K^+^ and to +35 mV in 100 mM K+, (40)) and in animals (HsKcnh1 Ea50 close to 0 mV in 5 mM K^+^; (26)). Finally, the high selectivity of RiSKC3 in favor of K^+^ with respect to Na^+^ (Fig. 7D) can contribute to the overall preferential accumulation of the former, over the latter in mycorrhizal roots, and thus to the increased tolerance to salt stress and plant growth under saline conditions. In conclusion, the present report provides evidence that rice endomycorrhizal symbiosis with *R. irregularis* improves the plant K^+^ nutrition and thereby tolerance to salt stress conditions, and that the Shaker voltage-sensitive K^+^ channel RiSKC3, dedicated to selective K^+^ secretion, appears as a member from the set of fungal K^+^ transport systems contributing to the mycorrhizal K^+^ transport pathway that ensures efficient and selective K^+^ translocation towards the host roots.

## Materials and Methods

### Plant Materials and Growth Conditions

Rice (*Oryza sativa* L.) ssp. Japonica cv. Kitaake plants were used in this study. *Rhizophagus irregularis* DAOM197198 (PremierTech, France) were used in this study. Seeds were germinated in a greenhouse (conditions: 16 h of day cycle, and 8 h of night at 28°C, 70% relative humidity) for a week. Germinated seedlings were then transplanted to sterilized quartz sand in 6 cm × 6 cm × 7.5 cm planting pots inoculated or not with the endomycorrhizal fungus, each pot receiving a single plant. For inoculation, a suspension of ∼500 spores of *R. irregularis* was added per pot.

The plants were submitted to two different treatments, qualified of either “high K^+^” or “low K^+^” condition, by irrigating the plants with Yoshida’s modified-solution (47) containing either 0.8 mM or 0.05 mM K^+^, respectively. The high K^+^ solution contained 0.7 mM KNO_3_, 0.05 mM K_2_SO_4_, 0.5 mM (NH_4_)_2_SO_4_, 1.6 mM MgSO_4_, 1.2 mM Ca(NO_3_)_2_, 60 µM FeSO_4_, 60 µM Na_2_EDTA, 20 µM MnSO_4_, 0.32 µM (NH_4_)_6_Mo_7_O_24_, 1.4 µM ZnSO_4_, 1.6 µM CuSO_4_, 45.2 µM H_3_BO_3_ and 0.25 mM NaH_2_PO_4_, pH 5-5.5. In the low K^+^ solution, 0.05 mM KNO_3_ was the single source of K^+^. The concentration of NO_3-_ was adjusted to 0.7 mM (as in the high K^+^ solution) by adding 0.65 mM NaNO_3_. The concentrations of the other salts were the same as in the high K^+^ solution, and the pH was adjusted to the same value.

### Salt treatment

Nine-week-old inoculated or non-inoculated rice plants (8 weeks post inoculation for the former plants) watered with Yoshida solution containing either 0.8 or 0.05 mM K^+^ were subjected to salt stress by adding 30 mL of the same Yoshida solution but supplemented with NaCl (100 mM) and Rb^+^, used as a tracer of K^+^ and provided (as RbCl) at the same concentration as that of K^+^ in the corresponding Yoshida solution, under conditions that prevented any leaching of the saline solution from the pot. Then, for one week, the plants were watered again with the initial Yoshida solution (free from added NaCl and RbCl), the solution being supplied in controlled amounts in order to avoid any leaching from the pot and thereby to maintain the salt stress conditions. Following this week of salt stress, the plants (then 10-week-old) were harvested and the shoot contents in K^+^, Rb^+^ and Na^+^ were determined.

### Measurement of K^+^, Rb^+^ and Na^+^ shoot contents

The collected shoots were weighed to determine their dry weight (DW). Ions were extracted from the tissues in 0.1 N HCl for 3 days and assayed by flame spectrophotometry (SpectrAA 220FS, Varian) (48).

### RNA extraction and transcript analysis

Total RNAs were extracted from roots using the RNeasy plus mini kit with gDNA eliminator according to manufacturer’s instructions (Qiagen, Germany). First-strand cDNAs were synthesized from 3 µg of RNAs using SuperScript III reverse transcriptase (Invitrogen) according to manufacturer’s instructions and used as template for RT-PCR or qRT-PCR experiments. PCR was performed as follows: 1 µL of cDNA (∼50 ng), 10 µL of 5× Green GoTaq® Reaction Buffer (Promega), 1 µL of dNTP Mix (10 mM each of dTTP, dCTP, dGTP, dATP), 5 µL of each of forward and reverse primers (10 µM, Supplemental Table 2), 0.25 µL of GoTaq® DNA Polymerase (5 U/µL, Promega), and 27.75 µL of ultra-pure water. PCR cycles (95°C for 2 min, and 30 cycles at 95°C for 30 sec, 60°C for 30 sec, 72°C for 30 sec and 72°C for 5 min) were performed in a FlexCycler^2^ (Analytikjena Biometra GmbH, Germany). The resulting PCR products were analysed by electrophoresis, stained with ClearSight DNA stain (Euromedex, France), and gel was scanned with Gel Doc^TM^ EZ Imager (Biorad). qRT-PCR analyses were performed using the Lightcycler480 system (Roche diagnostics) and SYBR *Premix Ex Taq* (Takara) in a total volume of 10 µL, which contained 2 µL of cDNA, 3 µL of forward and reverse primer mixture (1 µM, Supplemental Table 2), and 5 µL of SYBR *Premix Ex Taq*. Reactions were performed with three independent biological replicates, each one with three technical replicates (PCR program: 95°C for 30 sec; 45 cycles of 95°C for 10 sec, 60°C for 10 sec, and 72°C for 15 sec; followed by a melt cycle from 60°C to 95°C). CT (cycle threshold) values were obtained from amplification data using a threshold of 0.37.

### Liquid-phase *in situ* RT-PCR in rice roots

The protocol was modified from (49). Roots were cut into 1 cm small pieces and immediately fixed in PAA [2% (v/v) paraformaldehyde, 63% (v/v) ethanol, 5% (v/v) acetic acid] overnight at 4°C. Fixed root pieces were washed three times for 10 min each in DEPC-treated water. For reverse transcription, sections were incubated at 65°C for 5 min in a tube containing 2 μL of oligo(dT)_20_ (50 μM), 2 μL of 10 mM dNTP Mix (10 mM each dATP, dGTP, dCTP and dTTP) and 20 µL of ultra-pure water. Then the tubes were transferred into ice and added with 8 μL 5X First-Strand Buffer of SuperScript™ III RT kit (Invitrogen), 2 μL of 0.1 M DTT, 2 μL of RNaseOUT™ Recombinant RNase Inhibitor (40 units/μL, Invitrogen) and 2 μL of SuperScript™ III RT (200 units/μL, Invitrogen). The tubes were then incubated at 42°C for 1 h. The sections were washed once with DEPC-treated water. For PCR reaction, 10 µL of 5× Colorless GoTaq® Reaction Buffer (Promega), 1 µL of dNTP Mix (10 mM each of dTTP, dCTP, dGTP, dATP), 0.5 µL of digoxigenin-11-dUTP (1 nmol/µL, La Roche, Meylan, France), 5 µL of each of forward and reverse primers (10 µM, Supplemental Table S2), 0.25 µL of GoTaq^®^ DNA Polymerase (5 U/µL, Promega) and 28.25 µL of ultra-pure water were added into the tubes. PCR cycles (95°C for 2 min, and 30 cycles at 95°C for 30 sec, 60°C for 30 sec, 72°C for 30 sec and 72°C for 5 min) were performed in a FlexCycler^2^ (Analytikjena Biometra GmbH, Germany). Following RT-PCR, the tubes were washed three times for 10 min in 1× PBS (5 mM Na_2_HPO_4_, 130 mM NaCl, pH 7.5). After blocking with 2% (w/v) BSA in 1× PBS for 1 h, sections were incubated with alkaline phosphatase-conjugated anti-digoxigenin-Fab fragment (La Roche) diluted 1:250 in blocking solution for 2 h. The sections were rinsed three times with 10× washing buffer (100 mM Tris–HCl, 150 mM NaCl, pH 9.5) for 15 min each. Detection of alkaline phosphatase was carried out for ∼1 h using NBT/BCIP ready-to-use stock solution (La Roche) diluted in 1× washing buffer.

### Plasmid construction for expression of the RiSKC3-GFP fusion

A pFL61 plasmid (Genbank X62909) harbouring the construct *HcTOK1-GFP* under control of the *Saccharomyces cerevisiae phosphoglycerate kinase* (*PGK*) promoter and named pFL61[HcTOK1-GFP] was obtained from (46). The *HcTOK1* sequence was removed from this initial construct by PCR, the reaction medium containing 1 µL of pFL61[HcTOK1-GFP] (∼100 ng), 10 µL of 5× Phusion^®^ HF Buffer (Thermoscientific), 1 µL of dNTP Mix (10 mM each of dTTP, dCTP, dGTP, dATP), 5 µL of each of *UAL* forward and *LAPGF* reverse primers (10 µM, Supplemental Table S2), 0.5 µL of Phusion® DNA Polymerase (Thermoscientific) and 27.5 µL of ultra-pure water. PCR cycles (95°C for 2 min, and 25 cycles at 95°C for 30 sec, 60°C for 30 sec, 72°C for 3 min, and 72°C for 5 min) were performed in a FlexCycler^2^ (Analytikjena Biometra GmbH, Germany). The resulting construct was named pFL61[GFP]. A *RiSKC3* cDNA fragment was amplified by PCR using 1 µL of *RiSKC3* coding sequence cloned into a pGEM vector (∼100 ng), 10 µL of 5× Phusion® HF Buffer (Thermoscientific), 1 µL of dNTP Mix (10 mM each of dTTP, dCTP, dGTP, dATP), 5 µL of each of *UA846* forward and *LL846* reverse primers (10 µM, Supplemental Table S2), 0.5 µL of Phusion^®^ DNA Polymerase (2 U/µL, Thermoscientific) and 27.5 µL of ultra-pure water. PCR cycles (95°C for 2 min, and 25 cycles at 95°C for 30 sec, 60°C for 30 sec, 72°C for 1 min and 72°C for 5 min) were performed in a FlexCycler^2^ (Analytikjena Biometra GmbH, Germany). The resulting PCR products were analysed by electrophoresis, stained with ClearSight DNA stain (Euromedex, France), and the gel was scanned with Gel Doc^TM^ EZ Imager (Biorad). DNA fragments were excised from the gel by means of x-tracta™ Gel Extractor (Promega) and purified using Wizard® SV Gel and PCR Clean-Up System (Promega). Next, the In-Fusion HD Cloning Kit (Clontech, Mountain View, CA, USA) was used for cloning *RiSKC3* into pFL61[GFP] plasmid. The reaction medium contained 2 µL of 5× In-Fusion HD Enzyme Premix (Clontech); 1 µL of linearized pFL61[GFP] (∼100 ng), 1 µL of *RiSKC3* fragment (∼50ng) and 6 µL of ultra-pure water. The tube was incubated for 15 min at 50°C and placed on ice. One µL of the reaction medium was then used to transform Stellar^TM^ competent cells (Clontech). DNA plasmids were sequenced using the service of Eurofins Genomics (Germany).

### Heterologous expression in yeast

*Saccharomyces cerevisiae* strain PLY232 was used for heterologous expression of *RiSKC3-GFP* construct. Plasmid pFL61 harbouring the construct *RiSKC3-GFP* under the control of the *PGK* promoter was purified from Stellar^TM^ bacteria using the PureYield™ Plasmid Miniprep System (Promega). The yeast transformation protocol was described by (50).

### Confocal microscopy

A laser scanning confocal microscope (Leica TSC SP8 system, Germany) was used with the excitation wavelengths 488 nm and 561 nm for GFP and FM4-64, respectively. The detection wavelengths were in the range of 500-535 nm for GFP and 580-630 nm for FM4-64.

### Two-Electrode Voltage Clamp in Xenopus Oocytes

*RiSKC3* coding sequence was cloned into a modified pGEM-HE vector (D. Becker, University of Würzburg, Germany). Oocyte preparation, injection and two-electrode voltage-clamp (TEVC) recordings were performed as described (51). *In vitro* transcription was performed using the HiScribe T7 ARCA mRNA (with tailing) kit (New England Biolab, http://www.NEB.com) and Xenopus oocytes were injected with 30 ng of *RiSKC3* cRNA using a pneumatic injector. Oocytes were kept for 48 h at 20°C in a solution containing 96 mM NaCl, 2 mM KCl, 1.8 mM CaCl_2_, 2 mM MgCl_2_, 2.5 mM Na-pyruvate, 5 mM Hepes-NaOH (pH 7.5) and 50 µg/mL of gentamycin before recordings. Whole-cell currents were recorded using TEVC technique with oocytes bathed in K100 medium (100 mM KCl, 2 mM MgCl_2_, 1 mM CaCl_2_ and 10 mM HEPES, pH 7.5). Two second-long voltage steps to values ranging between –140 mV and +100 mV in +15 mV increments were applied from a −80 mV holding potential. For Erev determination, a tail-current protocol was used: after activation of RiSKC3 at +55 mV, voltage pulses were performed at voltages flanking Erev. Data acquisition and analyses were performed as previously described (52). All currents were obtained by subtraction of leak current at negative potentials.

### Phylogenetic tree construction

The protein sequences of the selected K^+^ channels from *Rhizophagus irregularis* (taxid:747089), *Hebeloma cylindrosporum* (taxid:76867), *Laccaria bicolor* (taxid:29883), *Physcomitrella patens* (taxid:3218), *Arabidopsis thaliana* (taxid:3702), *Drosophila melanogaster* (taxid:7227) and *Homo sapiens* (taxid:9606) were collected from databases by BLASTP analysis based on RiSKC3 (Supplemental Table S1). They were submitted to MUSCLE software for multiple alignment (53) The phylogenetic tree was generated with PhyML software (http://www.phylogeny.fr) using the maximum likelihood method (1000 bootstraps), and the tree was visualized using iTOL software (https://itol.embl.de/).

### Statistical Analysis

The data were analyzed by Student’s *t*-test to test differences between treatments.

## Acknowledgments

This project (ID 2204-001) was funded through LabEx AGRO ANR-10-LABX-0001-01 (under I-Site Université de Montpellier framework). This project was also supported by a Junior Research Fellowship Program from France Embassy in Thailand awarded to Dr Kawiporn Chinachanta. Confocal observations were performed at the Montpellier RIO Imaging facilities.

## Figures and Tables

**Supplemental Figure 1.**
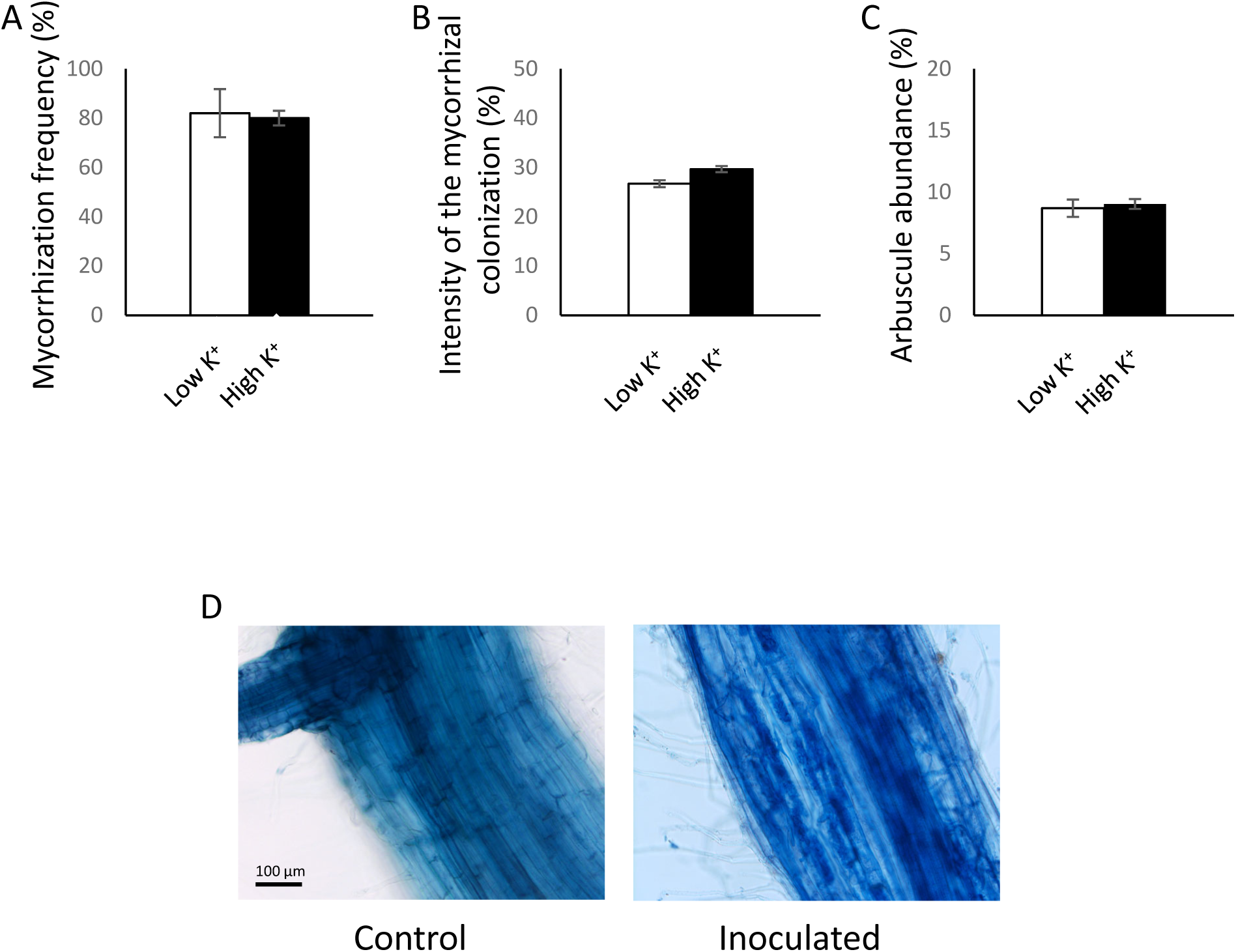
Estimation of AM colonization. After Trypan blue staining, roots were observed and mycorrhization frequency (A), intensity of the mycorrhizal colonization (B), and arbuscule abundance (C) were estimated using Mycocalc software (https://www2.dijon.inrae.fr/mychintec/Mycocalc-prg/download.html). Typical images of control and inoculated roots are shown (D).

**Supplemental Figure 2.**
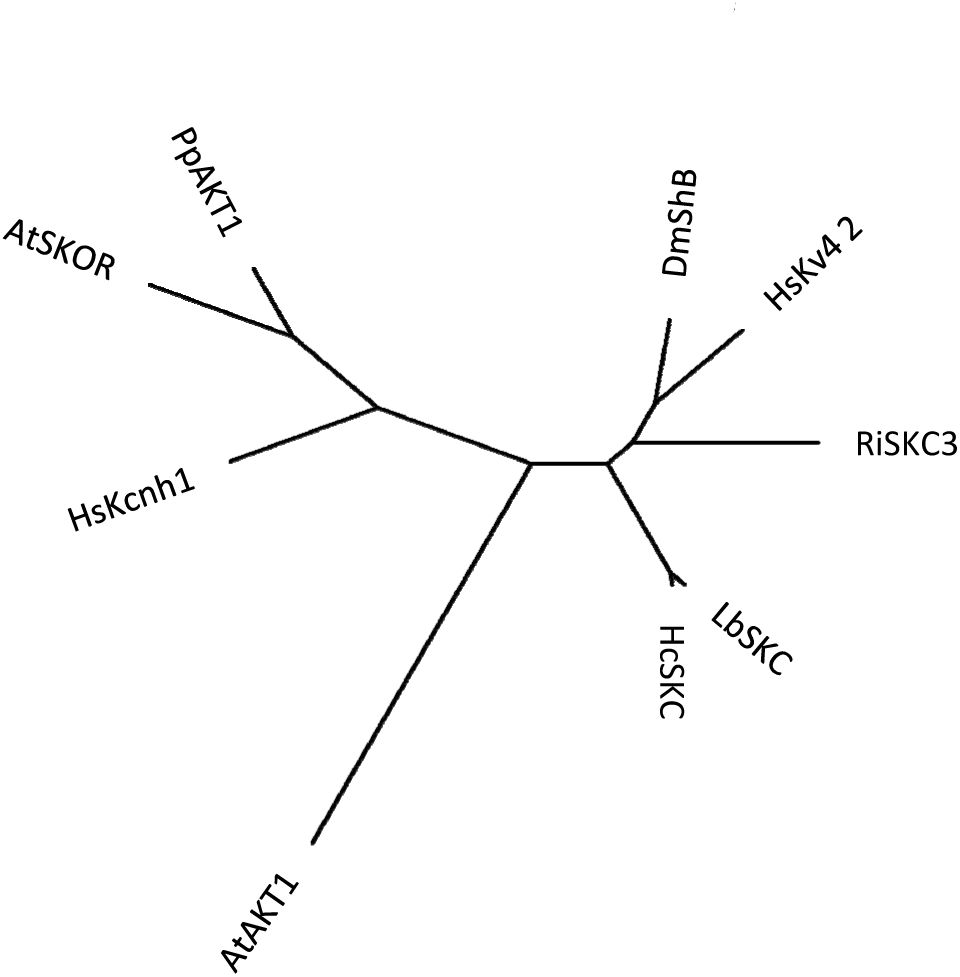
Phylogenetic analysis of Shaker channels in *R. irregularis*. The phylogenetic tree was generated with PhyML software (http://www.phylogeny.fr) using the maximum likelihood method. Branch length is proportional to the number of substitutions per site. RiSKC3 sequence was compared to the following sequences: AtAKT1 and AtSKOR from *Arabidopsis thaliana* (taxid: 3702,), DmShb from *Drosophila melanogaster* (taxid: 7227), HsKcnh1 and HsKv4,2 from *Homo sapiens* (taxid: 9606), PpAKT1 from *Physcomitrella patens* (taxid: 3218), HcSKC from *Hebeloma cylindrosporum* (taxid: 76867) and LbSKC from *Laccaria bicolor* (taxid: 29883). The protein (GenBank) accession numbers are given in Supplemental Table S1.

**Supplemental Figure 3.**
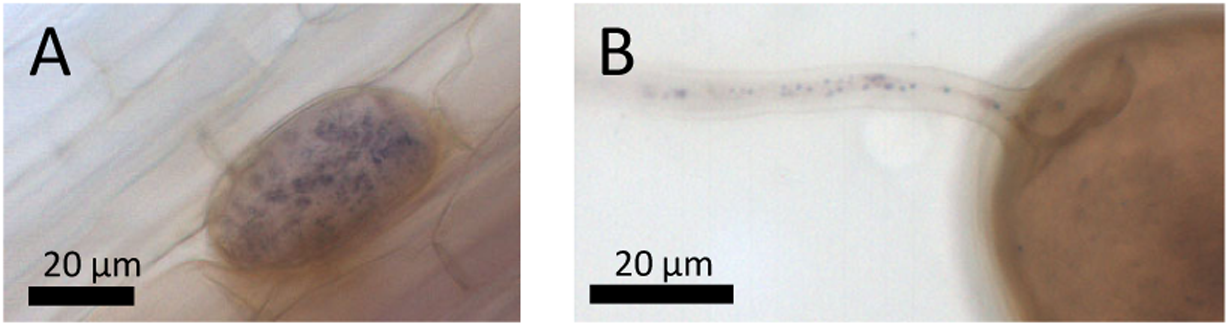
Endogenous phosphatase-alkaline activity in AM rice roots. Nine-week old roots of rice plants inoculated with *R. irregularis* were subjected to whole-mount *in situ* RT-PCR. Fixed tissues were incubated with NBT/BCIP to detect endogenous phosphatase-alkaline activity in vesicles (**A**) and in hyphae emerging from a spore (**B**) as a purple signal.

**Supplemental Table 1.** GenBank accession numbers of Shaker polypeptides present in phylogenetic trees.

| Gene name | Accession number | Protein ID |
| --- | --- | --- |
| RiSKC3 | GLOIN_2v1661846 | A0A015J981 |
| HcSKC |  | A0A0C3BDJ2 |
| LbSKC |  | B0DAX4 |
| PpAKT1 |  | A5PH36 |
| AtAKT1 |  | O22397 |
| AtSKOR |  | Q9M8S6 |
| DmShab |  | P08510 |
| HsKcnh1 |  | O95259 |
| HsKv4.2 |  | Q9NZV8 |

**Supplemental Table 2.**
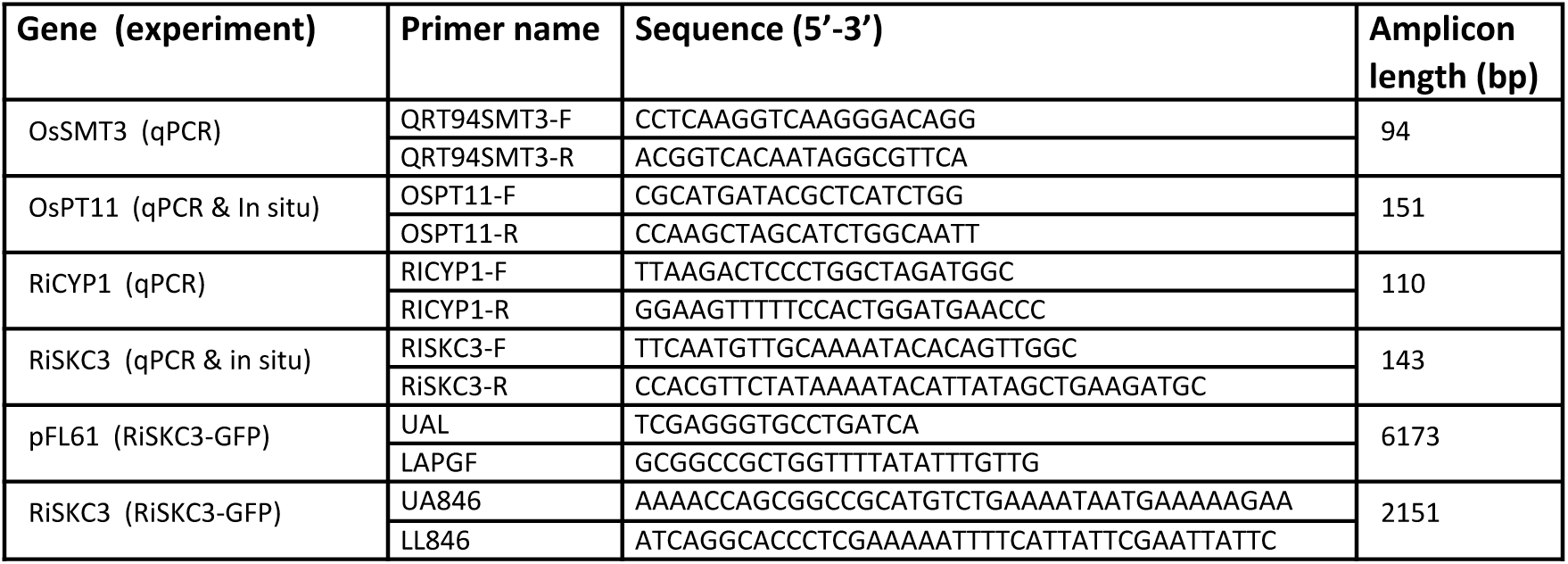
Primers used for PCR.

